# Within-host genetic micro-diversity of *Mycobacterium tuberculosis* and the link with tuberculosis disease features

**DOI:** 10.1101/2021.04.07.438754

**Authors:** Charlotte Genestet, Elisabeth Hodille, François Massol, Guislaine Refrégier, Alexia Barbry, Emilie Westeel, Gérard Lina, Florence Ader, Laurent Jacob, Stéphane Dray, Jean-Luc Berland, Samuel Venner, Oana Dumitrescu, on behalf of the Lyon TB study group

## Abstract

Tuberculosis (TB), caused by *Mycobacterium tuberculosis* (Mtb) complex, is still the number one deadly contagious disease. Mtb infection results in a wide spectrum of clinical presentations and severity symptoms, but without proven Mtb genetic determinants. Thanks to a collection of 355 clinical isolates with associated patient’s clinical data, we showed that Mtb micro-diversity within patient isolates is strongly correlated with TB-associated severity scores. Interestingly, this diversity is driven by a selection pressure to adapt to different lifestyles related to the infection site. Taken together, these results provide a new insight to better understand TB pathophysiology. Furthermore, Mtb micro-diversity could be envisioned as a new prognostic tool to improve the management of TB patients.

## INTRODUCTION

Tuberculosis (TB) caused by *Mycobacterium tuberculosis* (Mtb) complex remains one of the most prevalent and deadly infectious diseases, responsible for 10 million new cases and 1.2 million deaths among HIV-negative people worldwide in 2018, and an additional 208 000 deaths among HIV-positive people ^1^. Mtb infections result in a wide spectrum of clinical outcomes, from latent asymptomatic infection to pulmonary or extra-pulmonary manifestations of disease, with an array of severity symptoms. Such diversity has been historically attributed to host and environmental factors, while the Mtb complex was previously considered genetically monomorphic ^2^.

Since the introduction of the next generation sequencing (NGS) enabling whole genome sequencing (WGS), outstanding progress has been done in the field of Mtb genomics. Increasing studies based on NGS have revealed micro-diversity in Mtb clinical isolates: within hosts, minor variants coexist rather than a clonal colony ^3–14^. Although we showed in a previous study the involvement of initial Mtb micro-diversity in intra-macrophagic persistence and antibiotic tolerance ^15^, the role and the impact of Mtb micro-diversity is still poorly understood. It was already showed that mixed-strains Mtb infection is associated with poor outcome ^16,17^, but only few reports investigated the link between Mtb micro-diversity and TB severity and outcome ^9,10^. Yet, several major questions remain unanswered: such as the Mtb ability to cause active disease and various symptoms. While many Mtb virulence factors are well described, to date, there are no proven genetic determinants associated with virulence, disease progression or severity of TB ^18^.

Regarding other bacterial species responsible of chronic infections, such as *Helicobacter pylori, Pseudomonas aeruginosa, Staphylococcus aureus, Streptococcus pneumoniae*, several reports highlighted the role of micro-diversity in pathogen adaptation between and within-host. Bacterial micro-diversity plays a role in the adaptation to immune and treatment pressure, for infecting different body sites and has been suggested to impact outcome and severity of illness^19–25^.

Accordingly, here we explored the relation between Mtb micro-diversity of 355 clinical isolates, from 311 patients diagnosed at the Lyon University Hospital, and TB clinical presentation, as well as nutritional and immune status of patients. It revealed a strong correlation between the detection of Mtb micro-diversity within clinical isolates and TB-associated severity markers. Furthermore, thanks to a cohort of 42 patients with both microbiologically proven pulmonary and extra-pulmonary TB, we investigated intra-host micro-evolution of Mtb. We observed a compartmentalization of variants, driven by a selection pressure to adapt to different tissues, as shown by dN/dS approach. It should be noted that, besides the detection of canonical drug resistance determinants, the detection of Mtb micro-diversity within patient’s isolates is, to date, the only other bacterial feature which could be envisioned as a prognostic marker of poor TB outcome.

## RESULTS

### Baseline characteristics of study populations

A total of 355 clinical Mtb isolates from 311 patients were included in this study, including 42 patients with both microbiologically proven pulmonary and extra-pulmonary TB (PTB and EPTB) and 2 patients with 2 extra-pulmonary localization without PTB. Moreover, 25 patients with PTB also developed EPTB according to the clinical manifestations and 2 patients with EPTB may also have PTB but no pulmonary sampling was conducted. The **Table 1** summarize patients’ demographic and descriptive characteristics. As it is a retrospective study, all data were not available for all patients, then the population size was always specified.

**Table 1:** Patients’ demographic and baseline descriptive characteristics. Data were expressed as count (percentage, %) for dichotomous variables and as medians (interquartile range [IQR]) for continuous values. The number of missing values was excluded from the denominator. Fisher exact, χ2 test or non-parametric Mann-Whitney U test was used to compare groups where appropriate. *p-value* < 0.05 was considered significant. PTB: pulmonary tuberculosis; EPTB: extrapulmonary tuberculosis; unkown outcome: loss of follow-up or follow-up in another care facility; HIV: human immunodeficiency virus; MUST: malnutrition universal screening tool; BMI: body mass index; HDL-C: high-density lipoprotein cholesterol; LDL-C: low-density lipoprotein cholesterol; CRP: C-reactive protein; WBC: white blood cell.

| Characteristics | Pulmonary TB<br>n=244 | Extra-pulmonary TB<br>n=111 | *p-values* |
| --- | --- | --- | --- |
| **Demographic data** |  |  |  |
| Age (years), median [IQR] | 36 [25-58] | 37 [24-60] | 0.7505 |
| Gender (male), n (%) | 162 (66.4) | 57 (51.4) | **0.0094** |
| **TB outcome** |  |  |  |
| Fatal outcome, n (%) | 8 (3.3) | 3 (2.7) | 0.6999 |
| Unknown, n (%) | 18 (7.4) | 11 (9.9) | 0.4109 |
| **Comorbidity** |  |  |  |
| HIV, n (%) | 15/233 (6.4) | 8/107 (7.5) | 0.8165 |
| Hepatitis, n (%) | 24/189 (12.7) | 13/85 (15.3) | 0.5701 |
| Diabetes, n (%) | 26/234 (11.1) | 14/105 (13.3) | 0.5867 |
| Immunosuppressive therapy, n (%) | 21/235 (8.9) | 16/107 (15.0) | 0.1317 |
| History of TB, n (%) | 19/233 (8.2) | 4/106 (3.8) | 0.1662 |
| **TB-associated severity scores** |  |  |  |
| Bandim TB score, median [IQR] (n) | 4 [3-6] (233) | 3 [1.5-5] (105) | **0.0002** |
| MUST, median [IQR] (n) | 3 [1-5] (227) | 2 [0-4] (105) | **0.0293** |
| Nutritional risk score, median [IQR] (n) | 1 [0.25-3] (56) | 1 [0-3] (24) | 0.8839 |
| **Nutritional status** |  |  |  |
| BMI (kg/m<sup>2</sup>), median [IQR] (n) | 20.2 [17.7-22.4] (228) | 21.63 [19.0-26.2] (107) | **<0.0001** |
| Weightloss, median [IQR] (n) | 8.0 [4.5-12.3] (227) | 6.4 [2.4-11.5] (106) | **0.0293** |
| Serum albumin (g/L), median [IQR] (n) | 30.0 [26.0-36.7] (209) | 31.1 [25.9-36.2] (92) | 0.8693 |
| Serum pre-albumin (g/L), median [IQR] (n) | 0.13 [0.09-0.18] (152) | 0.11 [0.08-0.17] (60) | 0.3448 |
| **Serum/blood parameters** |  |  |  |
| Total protein (g/L), median [IQR] (n) | 75 [70-81] (235) | 75 [70-81] (102) | 0.9509 |
| Blood glucose (mmol/L), median [IQR] (n) | 5.0 [4.5-6.0] (215) | 5.2 [4.6-6.4] (100) | 0.3719 |
| Calcium (mmol/L), median [IQR] (n) | 2.44 [2.31-2.61] (236) | 2.43 [2.32-2.59] (92) | 0.7641 |
| Phosphorus (mmol/L), median [IQR] (n) | 0.98 [0.85-1.18] (118) | 1.07 [0.84-1.21] (57) | 0.6397 |
| Magnesium (mmol/L), median [IQR] (n) | 0.81 [0.74-0.88] (70) | 0.80 [0.71-0.87] (42) | 0.2684 |
| Sodium (mmol/L), median [IQR] (n) | 137 [134-139] (236) | 137 [134-139] (62) | 0.1480 |
| Potassium (mmol/L), median [IQR] (n) | 4.0 [3.8-4.3] (236) | 4.0 [3.7-4.3] (104) | 0.3781 |
| Chloride (mmol/L), median [IQR] (n) | 103 [100-106] (235) | 103 [98-106] (103) | 0.2117 |
| Bicarbonate (mmol/L), median [IQR] (n) | 25 [23-27] (178) | 25 [23-27] (80) | 0.8603 |
| Anion gap, median [IQR] (n) | 13 [11-14] (165) | 13 [11-15] (78) | 0.3953 |
| **Serum lipid profile** |  |  |  |
| Total cholesterol (mmol/L), median [IQR] (n) | 4.1 [3.3-5.1] (56) | 4.2 [3.1-4.9] (24) | 1.0000 |
| HDL-C (mmol/L), median [IQR] (n) | 1.03 [0.69-1.29] (52) | 1.00 [0.54-1.19] (23) | 0.7302 |
| LDL-C (mmol/L), median [IQR] (n) | 2.58 [1.88-3.33] (50) | 2.58 [1.99-3.25] (23) | 0.8960 |
| Triglyceride (mmol/L), median [IQR] (n) | 1.14 [0.93-1.67] (59) | 1.24 [0.99-1.69] (28) | 0.4898 |
| **Iron balance and inflammatory markers** |  |  |  |
| Serum iron (μmol/L), median [IQR] (n) | 5.3 [3.0-9.2] (73) | 6.7 [4.5-10.6] (39) | 0.1555 |
| Transferrin saturation (%), median [IQR] (n) | 13.1 [8.2-20.1] (73) | 13.4 [10.1-20.3] (39) | 0.2921 |
| Ferritin (μg/L), median [IQR] (n) | 195 [79-319] (98) | 281 [104-759] (48) | **0.0336** |
| CRP (mg/L), median [IQR] (n) | 48 [13-100] (228) | 45 [14-93] (99) | 0.8781 |
| **Hemogram** |  |  |  |
| Hemoglobin (g/L), median [IQR] (n) | 120 [104-136] (236) | 115 [102-130] (106) | 0.1644 |
| WBC (G/L), median [IQR] (n) | 6.9 [5.3-9.0] (237) | 6.3 [4.6-8.1] (105) | **0.0294** |
| Neutrophils (G/L), median [IQR] (n) | 4.6 [3.2-6.2] (237) | 4.4 [2.8-5.5] (105) | 0.1218 |
| Eosinophils (G/L), median [IQR] (n) | 0.09 [0.03-0.18] (236) | 0.075 [0.01-0.148] (104) | **0.0446** |
| Basophils (G/L), median [IQR] (n) | 0.03 [0.02-0.05] (235) | 0.02 [0.01-0.04] (104) | 0.0591 |
| Monocytes (G/L), median [IQR] (n) | 0.62 [0.46-0.84] (237) | 0.55 [0.38-0.75] (105) | **0.0179** |
| Lymphocytes (G/L), median [IQR] (n) | 1.41 [0.89-2.00] (237) | 1.18 [0.75-1.68] (105) | **0.0294** |
| **Lymphocyte typing** |  |  |  |
| CD4 T cells (cell/mm<sup>3</sup>), median [IQR] (n) | 338 [180-613] (81) | 362 [198-555] (45) | 0.8406 |
| CD8 T cells (cell/mm<sup>3</sup>), median [IQR] (n) | 274 [168-468] (79) | 240 [151-384] (44) | 0.1581 |
| CD4/CD8 ratio, median [IQR] (n) | 1.43 [0.82-2.07] (79) | 1.77 [0.84-2.5] (44) | 0.2249 |

Patient’s descriptive characteristics revealed different profiles between patients with PTB or EPTB. Overall, higher severity scores and lower malnutrition indices were observed for patient with PTB compared to patients with EPTB. Conversely, a lower immune status was observed for patient with EPTB compared to patients with PTB. It should be noted that serum lipid and serum iron profiles and lymphocyte typing can not be properly compared between the cohorts as the data were available only for 20% to 43% of patients.

### Mtb micro-diversity is not correlated with Mtb samples characteristics

WGS data of Mtb clinical isolates included in this study were analyzed to detect Mtb genetic micro-diversity through unfixed mutations (at frequencies between 10 and 90%). These data were used to identify the minimum number of variants in each Mtb clinical isolates. A variant was defined as an assembly of mutations with similar frequencies (±10%). This allowed to calculate the α-diversity of each Mtb isolates, which include both the number and the frequency of the variants and the genetic distance between these variants.

First of all, correlation between Mtb micro-diversity and Mtb samples characteristics, such as smear results and time to positivity of Mtb culture samples, reflecting bacterial load in clinical isolates, type of extra-pulmonary or pulmonary samples, Mtb lineages and resistance status, were explored (**Fig. S1**). Risks of bias, such as delta between Mtb sampling or between treatment initiation and sampling (only concerning the patients with both microbiologically proven PTB and EPTB) and samples that underwent freeze / thaw cycle (biobank samples) compared to those analyzed after a single round of culture (routine practice), were also evaluated (**Fig. S2**). No significant correlation was observed, except a higher proportion of isolates with captured micro-diversity in biobank pulmonary samples compared to pulmonary samples analyzed in routine practice. It may be due to the selection of significant Mtb isolates from our biobank, such as MDR strains, highly transmissible Mtb strains and strains from patients with both PTB and EPTB, all factors usually associated with high TB severity. Interestingly, no significant correlation was established between the bacterial load and Mtb micro-diversity in clinical isolates.

### Mtb micro-diversity is correlated with TB-associated severity indices

We aimed to investigate the association between Mtb α-diversity and TB outcome. However, due to the very low rate of fatal outcome in our setting (8/311; 2.6%), no significant correlation could be established (**Fig. S3**). Therefore, the correlation between Mtb micro-diversity and 3 TB severity indices was explored: i) the modified Bandim score (the most largely used PTB prognosis score), which consider 5 symptoms (cough, hemoptysis, dyspnea, chest pain, night sweats) and 5 clinical findings (anemia, tachycardia, positive finding at lung auscultation, fever, BMI) ^26,27^; ii) the Malnutrition Universal Screening Tool (MUST), based on weight loss, BMI and anorexia to evaluate malnutrition status of TB patients ^28^; iii) the nutritional risk score considering BMI, hypoalbuminemia, hypocholesterolemia and severe lymphocytopenia, then including both nutritional and immune factors ^29^.

A significant correlation was observed between the detection of micro-diversity (α-diversity > 1) in pulmonary Mtb isolates and high Bandim score (**Fig. 1A**, *p*<0.0001), MUST (**Fig. 1B**, *p*<0.0001) and nutritional risk score (**Fig. 1C**, *p*<0.0001). However, no correlation was observed between the ranges of α-diversity in Mtb isolates and the TB-associated severity indices evaluated.

**Figure 1:**
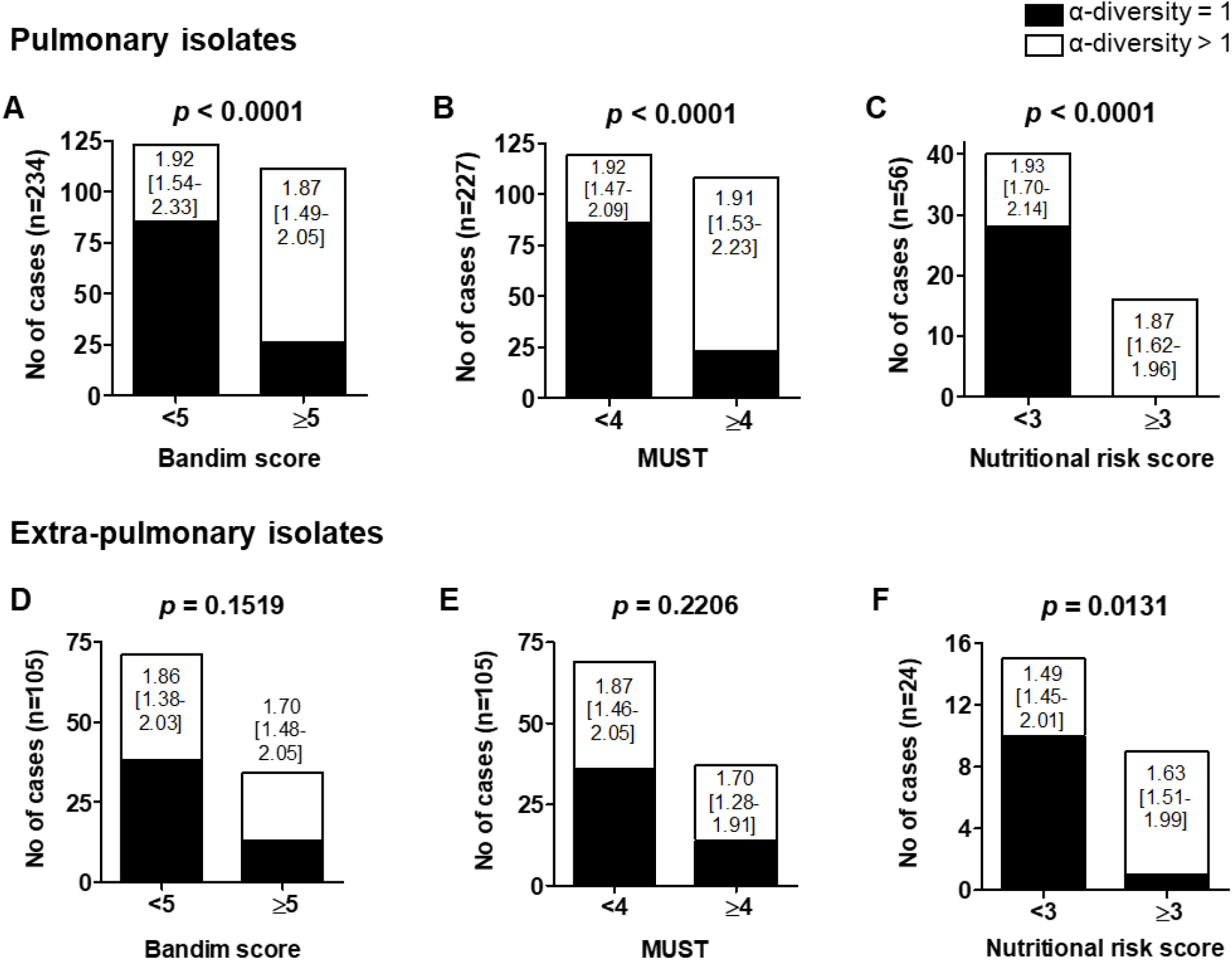
The detection of genetic micro-diversity in Mtb isolates is associated with TB-associated severity indices. Association between the detection and the range of Mtb α-diversity from pulmonary (A-C) or extra-pulmonary samples (D-F) and TB-associated severity indices, the Bandim score (A, D), the MUST (malnutrition universal screening tool, B and E), and the nutritional risk score (C, F). Black bar: α-diversity=1 no diversity detected by WGS; white bar: α-diversity>1 at least two variants detected by WGS. *p* = x.xxxx: non-parametric statistical method Fisher exact test was used to compare groups. *p-value* < 0.05 was considered significant. x.xx [y.yy-z.zz]: median [IQR] of Mtb α-diversity. Mann-Whitney U test was used to compare ranges of Mtb α-diversity between groups (no statistical differences observed).

As expected, no correlation was observed between the detection of micro-diversity in extra-pulmonary Mtb isolates and the Bandim score (**Fig. 1D**), as this score is mainly based on PTB symptoms, nor with the MUST (**Fig. 1E**), as malnutrition is more characteristic of PTB (**Table 1**) 30. Conversely, a correlation between the detection of micro-diversity in extra-pulmonary Mtb isolates and high nutritional risk score (**Fig. 1F**, *p*=0.0131) was observed, this severity index also including variables of the patient’s immune status. As before, no correlation was observed between the ranges of α-diversity in Mtb isolates and the TB severity indices evaluated.

### Independent variables associated with micro-diversity in Mtb pulmonary and extra-pulmonary isolates

The analysis of nutritional indices of TB patients revealed that the detection of micro-diversity in pulmonary Mtb isolates was correlated with low BMI, severe unintentional weight loss, low serum pre-albumin, hyponatremia, hypochloremia, low serum bicarbonate (**Fig. S4**), and also with low total cholesterol, low HDL and high triglyceride (**Fig. S5**), which are all associated with unfavorable TB outcome ^29–34^. The detection of micro-diversity in extra-pulmonary Mtb isolates was correlated with severe unintentional weight loss, low serum pre-albumin and low total serum protein (**Figs. S5 and S6**). Furthermore, a correlation was observed between the detection of micro-diversity in both Mtb pulmonary and extra-pulmonary isolates and iron deficiency, meaning low serum iron and low transferrin saturation (**Fig. 2**).

**Figure 2:**
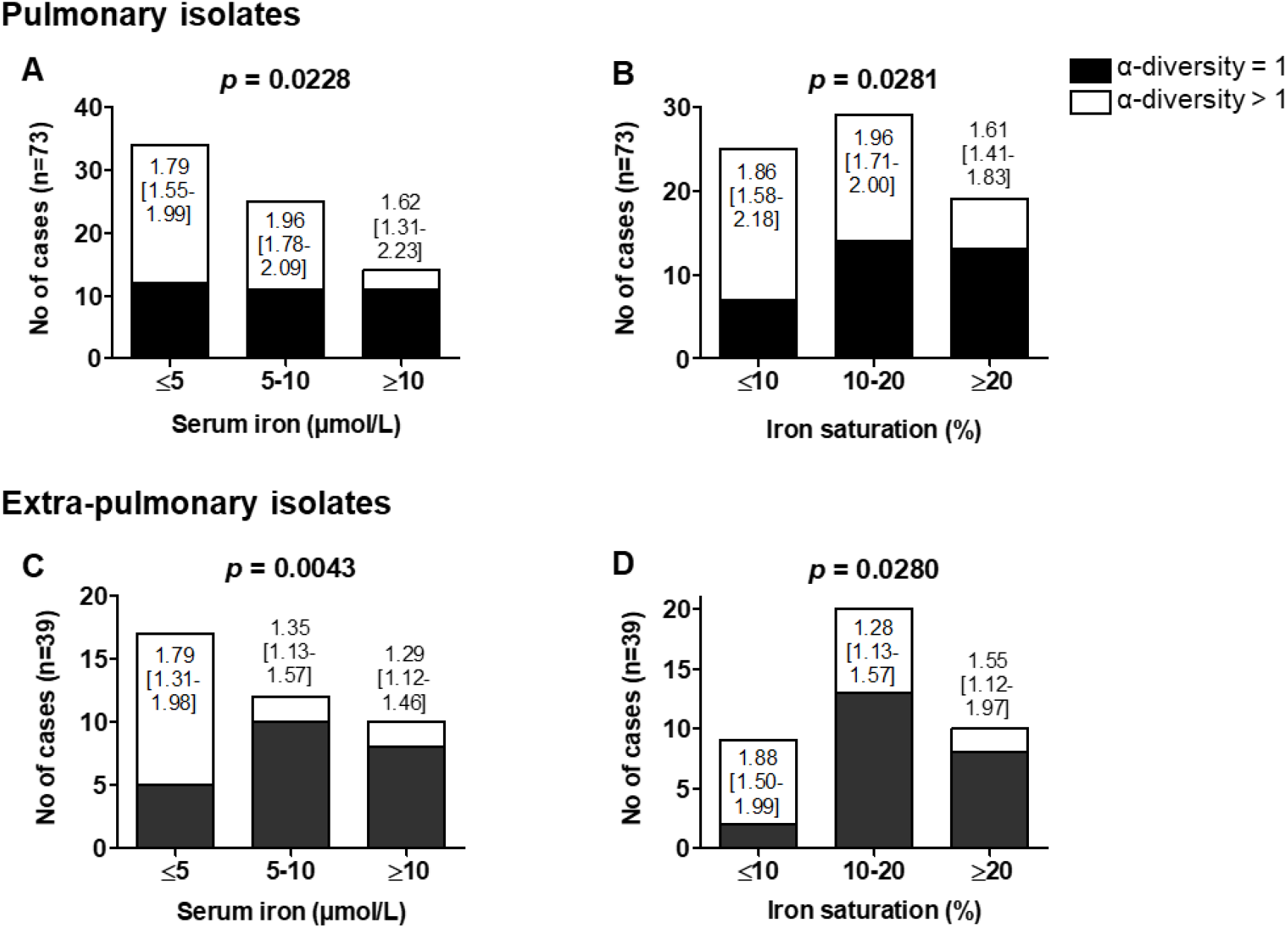
The detection of genetic micro-diversity in pulmonary and extra-pulmonary Mtb isolates is associated with iron deficiency. Association between the detection and the range of Mtb α-diversity from pulmonary (A and B) and extra-pulmonary (C and D) samples and serum iron level (A and C) and transferrin saturation (B and D). Black bar: α-diversity=1 no diversity detected by WGS; white bar: α-diversity>1 at least two variants detected by WGS. *p* = x.xxxx: non-parametric statistical method χ2 test was used to compare groups. *p-value* < 0.05 was considered significant. x.xx [y.yy-z.zz]: median [IQR] of α-diversity. Kruskal-Wallis analysis, using Dunn’s Multiple Comparison Test was used to compare ranges of α-diversity between groups (no statistical differences observed).

We also explored the correlation between micro-diversity in Mtb isolates and inflammation (serum CRP and ferritin) and immune markers (hemoglobin, blood white blood cell (WBC), neutrophil, eosinophil, basophil, monocyte, lymphocyte, CD4 T cells and CD8 T cells count in peripheral blood and CD4/CD8 ratio). No correlation was observed between these markers and the detection of micro-diversity in pulmonary Mtb isolates (**Fig. S7**). Conversely, we found a correlation between the detection of micro-diversity in extra-pulmonary Mtb isolates and high serum ferritin and low CD4 T cells count in peripheral blood (**Fig. S8**), which is a well-known risk factor for TB patients, especially for extra-pulmonary TB ^35,36^. Finally, in all cases (nutritional and immune variables), except for phosphorus for pulmonary Mtb isolates and eosinophils count for extra-pulmonary Mtb isolates, no correlation was observed between the ranges of Mtb α-diversity and the host variables studied.

### Models describing the relationship between Mtb micro-diversity and clinical parameters

To go forward, we built a model to relate Mtb micro-diversity in pulmonary (P) and extra pulmonary (EP) locations to various explanatory variables obtained for most patients in the cohort. The response variable we chose to model is a binary (0/1) measure of whether Mtb micro-diversity was detected in clinical isolates.

In the case of pulmonary TB infections, model comparisons indicated a best model with only 9 variables (double location of infection, Mtb lineage, weight loss, protein dosage, CRP, hemoglobin, leukocyte count, neutrophil count and the BANDIM score). 39 models had an Akaike weight less than two units from the best model. Only eight variables had likely effects (BANDIM score, Weight loss, Double Location, Hemoglobin, Neutrophil count, CRP, Protein, and Leukocyte count), and only the effect of Mtb lineage had importance between 0.5 and 0.73, making it a plausible (but not likely) effect (**Fig. 3**). Multi-model inference of parameter values ascertained that Mtb micro-diversity in pulmonary isolates was higher with increasing weight loss, BANDIM, hemoglobin, and leukocyte counts, and lower with increasing neutrophil counts, CRP, and protein (**Table 2**). All lineages had slightly different coefficients, but they never differed from the baseline sufficiently for 95% confidence intervals to have the same sign (**Table 2**) and asymptotic pairwise tests between lineage coefficients were all non-significant in the best model (results not shown).

**Figure 3:**
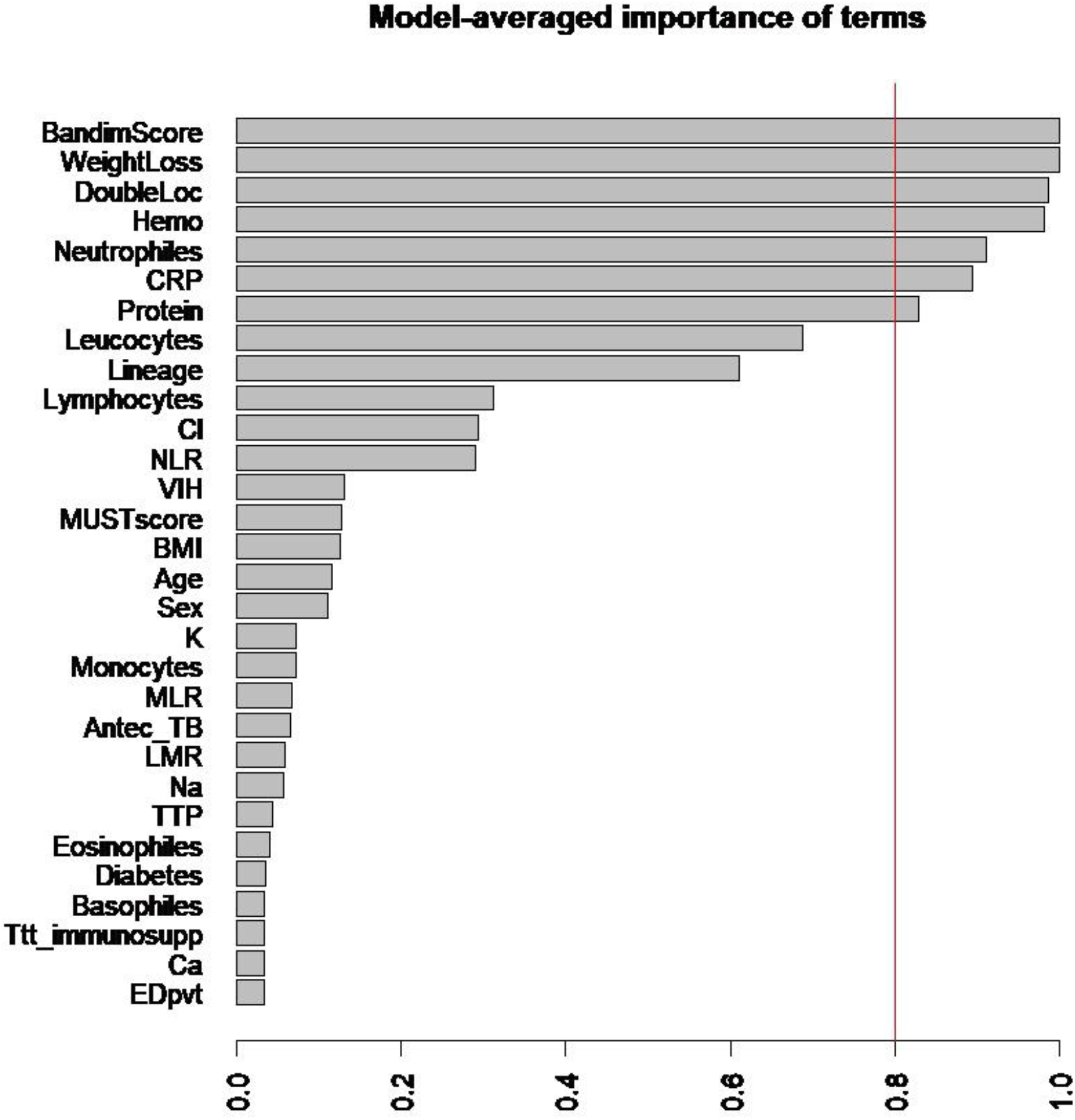
Model-averaged importance of terms for Mtb micro-diversity in pulmonary isolates. Model-averaged importance of each term in the model (**Table 2**), which is defined as the proportion of the 2000 best models in which a given term appears. Red line indicates 80% support. Terms with an importance above the red line are included in our final model.

**Table 2:** Models of Mtb micro-diversity in pulmonary isolates. Multi-model estimates of variable coefficients based on the 2000 models pooled in the consensus set of models; only models with importance greater than 0.5 were kept in this table. “DoubleLoc1”: presence of both P and EP locations of TB infection in the patient, based on microbiological or pathophysiological evidence; “Lineage2”: effect of lineage 2 when compared to baseline (lineage 1). “Nb models”: number of models in the consensus set (already the result of merging four parallel model comparison processes) which incorporated the variable.

|  | Estimate | lower CI | higher CI | Uncond. variance | Nb models | Importance |
| --- | --- | --- | --- | --- | --- | --- |
| (Intercept) | 2.36E-01 | -1.46E+01 | 1.50E+01 | 6.71E+01 | 2000 | 1.00 |
| WeightLoss | 2.73E-01 | 1.58E-01 | 3.89E-01 | 3.49E-03 | 2000 | 1.00 |
| BandimScore | 5.85E-01 | 3.15E-01 | 8.55E-01 | 1.88E-02 | 2000 | 1.00 |
| DoubleLoc1 | 1.26E+00 | 2.03E-01 | 2.31E+00 | 2.93E-01 | 1962 | 0.99 |
| Haemoglobin | 2.84E-02 | 4.43E-03 | 5.24E-02 | 1.52E-04 | 1957 | 0.99 |
| Neutrophiles | -6.36E-04 | -1.52E-03 | 2.51E-04 | 2.17E-07 | 1827 | 0.93 |
| CRP | -8.97E-03 | -1.89E-02 | 1.00E-03 | 2.70E-05 | 1772 | 0.92 |
| Protein | -4.81E-02 | -1.08E-01 | 1.20E-02 | 1.00E-03 | 1654 | 0.87 |
| Leucocytes | 4.06E-04 | -3.72E-04 | 1.18E-03 | 1.67E-07 | 1384 | 0.74 |
| Lineage2 | -6.54E-01 | -2.86E+00 | 1.55E+00 | 1.37E+00 | 1224 | 0.65 |
| Lineage3 | -2.11E+00 | -6.16E+00 | 1.95E+00 | 4.26E+00 | 1224 | 0.65 |
| Lineage4 | -6.93E-01 | -2.82E+00 | 1.44E+00 | 1.26E+00 | 1224 | 0.65 |
| Lineage6 | -1.60E-01 | -2.52E+00 | 2.20E+00 | 2.01E+00 | 1224 | 0.65 |
| LineageM. bovis | -3.13E+00 | -8.59E+00 | 2.32E+00 | 7.82E+00 | 1224 | 0.65 |

### Mtb micro-diversity within and between pulmonary and extra-pulmonary compartments

To better understand the role and the impact of Mtb micro-diversity, we focused on paired isolates from the training cohort to explore micro-evolution of Mtb within individuals (**Fig. 4**). First of all, the frequency in both pulmonary and extra-pulmonary compartments of each variant identified was explored. Among the 104 variants identified by WGS, 31/104 (30%) were specific of pulmonary isolates, 30/104 (29%) were specific of extra-pulmonary isolates, the frequencies of 22/104 (21%) variants were significantly increased or decreased (≥10%) between the compartments and only 21/104 (20%) were found at similar frequencies between the compartments (**Fig. 4A**). These results suggest a compartmentalization of Mtb variants.

**Figure 4:**
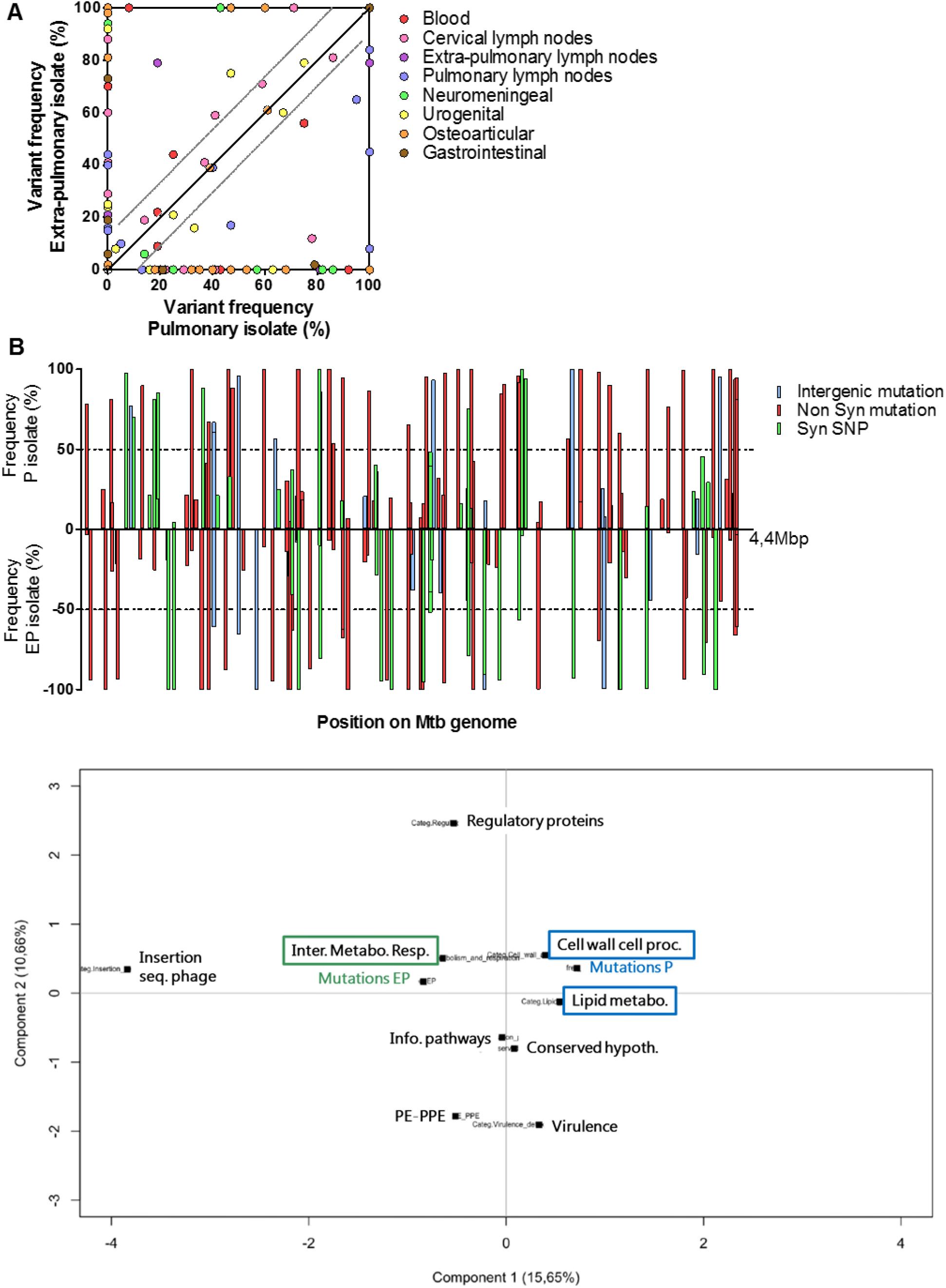
Compartmentalization of Mtb variants between pulmonary and extra-pulmonary compartments. (A) Frequencies in pulmonary and extra-pulmonary isolates of the 104 variants identified in paired isolates from the 42 patients of the training cohort with both microbiologically proven pulmonary and extra-pulmonary TB. (B) Repartition on Mtb reference genome of the 168 pairwise mutations distance identified between the 42 paired isolates from the training cohort. Each bar represents a pairwise mutation distance between paired pulmonary and extra-pulmonary isolates. *y-axis* positive value: mutation frequency in pulmonary isolate; *y-axis* negative value: mutation frequency in extra-pulmonary isolate. Bleu bar: intergenic mutation; red bar: nonsynonymous mutation; green bar: synonymous single nucleotide polymorphism (SNP). (C) Principal component analysis (PCA) for mixed data was performed with the delta of frequency of the 104 nonsynonymous pairwise mutation observed between paired pulmonary and extra-pulmonary isolates and the functional categories of these mutations. The first principal component allowed a discrimination of pulmonary and extra-pulmonary mutations.

Then we analyzed of the repartition on Mtb genome of the 168 pairwise mutations distances observed between paired pulmonary and extra-pulmonary isolates, among these 104 variants. However, it did not reveal any hot spot on Mtb genome (**Fig. 4B**). Among these 168 pairwise mutations, 22/168 (13%) were intergenic mutations, and 146/168 (87%) were located in coding regions, among whom 104/146 (71%) were nonsynonymous mutations and 42/146 (29%) were synonymous SNP.

These 104 nonsynonymous mutations were used to perform a principal component analysis (PCA) for mixed data (**Fig. 4C**). This analysis took into account the delta of frequency of each pairwise mutation between pulmonary and extra-pulmonary compartment and the functional categories of these mutations. The first principal component (15.65% of the variance) allowed a discrimination of pulmonary and extra-pulmonary mutations and of some functional categories. This revealed an association of pulmonary mutations with cell wall and cell processes and lipid metabolism categories. Otherwise, extra-pulmonary mutations were strongly associated with intermediary metabolism and respiration category. This suggests, on the one hand, an adaptation to pulmonary macrophage infection for pulmonary variants and on the other hand, a metabolic adaptation to extra-pulmonary tissues.

To go forward, we used PAML branch-site models to test for selection pressures associated with infection localization. No gene function exhibited a significant signal of positive selection pressure, neither in pulmonary isolates nor in extra-pulmonary ones with our cohort. Still, relatively low p-values (0.1<p-value <0.15) combined with amino acids with significant BEB posterior P-values were observed for “Virulence, detoxification and adaptation” gene category in pulmonary Mtb isolates and for “intermediary metabolism and respiration” gene category in the extra-pulmonary ones. These tendencies require larger samples to be confirmed.

## DISCUSSION

The objective of the present study was to address the impact of Mtb micro-diversity on TB pathophysiology. For this purpose, we explored the correlation between Mtb micro-diversity and TB clinical presentation, meaning pulmonary and extra-pulmonary tuberculosis and severity of illness. A strong correlation was observed between the detection of Mtb micro-diversity and TB-associated severity markers, in both pulmonary and extra-pulmonary clinical isolates. Furthermore, this diversity seems to be driven by a selection pressure to adapt to different tissues.

A previous report explored the correlation between baseline Mtb diversity (initiation or major change to treatment) and 6-month TB outcome and found that Mtb diversity did not affect TB outcome. However, beside fatal outcome, treatment failure and acquisition of multi-drug resistance, authors also included loss of follow-up in unfavorable outcome ^9^. Nevertheless, the latter category, representing 33% of poor outcome in the study, is not itself an unfavorable outcome, therefore we have classified it in the “unknown” category, which could explain the discrepancies observed between the studies. Otherwise, our results are in accordance with another study, focusing on the evolution of Mtb micro-diversity from five patients during the course of TB treatment, which showed that higher TB severity is associated with an increase of Mtb micro-diversity within-host, and more particularly in pre-mortem Mtb isolates of two of these patients, and without significant impact of TB treatment on Mtb micro-diversity ^10^. Yet it is still unclear whether Mtb micro-diversity is a cause (better Mtb adaptation to treatment, to immune pressure and/or to various niches) or a consequence (tissue breakdown allowing sampling of Mtb variants usually inaccessible and/or lower immune response reducing selection pressure) of the TB severity. The mechanisms driving such diversity remain to be explored. As sown using PAML branch-site models, variants harbored SNP in different functional categories according to their localization, meaning pulmonary or extra-pulmonary samples. Even if the tendencies observed require larger samples to be confirmed, the result obtained suggested an adaptation to pulmonary macrophage infection for pulmonary Mtb isolates and a metabolic adaptation for extra-pulmonary ones. Moreover, development of *in vitro* models will be needed to decipher the role and the impact of the identified pairwise variants between pulmonary and extra-pulmonary compartments.

Alongside that, in the present study, no correlation was found between the ranges of Mtb α-diversity and TB-associated severity markers. It may be due to the fact that the analysis was based on minimum number of variants estimated through WGS data to calculate Mtb α-diversity, which was a risk to underestimate micro-diversity in Mtb clinical isolate. However, to be exhaustive regarding the variant composition of an Mtb isolate, this would have required sequencing of several colonies for each Mtb clinical isolate, which is time- and cost-consuming. Nevertheless, detection of unfixed mutations at the level of WGS (meaning mutation frequencies between 10 and 90%) was enough to observe a strong correlation between Mtb micro-diversity detection and TB-associated severity markers.

As WGS is performed in routine practice in our lab, as well as in other TB diagnosis lab, it could be envisioned as an all-in-one solution, to detect antibiotic resistance ^37^, to infer Mtb transmission chains and to perform epidemiological monitoring ^38,39^ and now as a prognosis tool. In the frame of cancer and microbiological research, calling algorithm for low frequency variants were developed ^40–42^ and may be adapt to Mtb WGS data. Therapeutic drug monitoring and implementation of additional management measures could be performed for patients with detectable Mtb micro-diversity in clinical isolates. It would ensure optimal anti-TB drug doses and prevent slow response to treatment, which would reduce risks of treatment failure and of drug resistance acquisition ^43,44^.

In conclusion, although these results need to be confirmed in an independent prospective validation study, Mtb micro-diversity within clinical isolate could be a useful prognosis tool to ensure optimal management of TB patients.

## METHODS

### Ethical considerations

For this study we recorded demographical (age, sex), clinical (extrapulmonary and/or pulmonary TB), microbiological (smear sputum results, growth delay, antibiotic resistance, lineage) and nutritional and immune data. All data were implemented in a database, in accordance with the decision 20-216 of the ethics committee of the Lyon University Hospital, France and the French Bioethics laws (Reference methodology MR-004 that covers the processing of personal data for purposes of study, evaluation or research that does not involve the individual). Relevant approval regarding access to patient-identifiable information are granted by the French data protection agency (Commission Nationale de l’Informatique et des Libertés, CNIL).

### Mtb samples and data collection

In this retrospective study, a total of 355 Mtb clinical isolates were included, from 311 patients diagnosed with microbiologically proven TB at the Lyon University Hospital. We included all Mtb clinical isolates for which WGS was performed in routine practice at the Lyon University Hospital from January 2017 to January 2020 and significant isolates of our collection (MDR Mtb, representative strains of large previously identified clusters, samples from patients with both pulmonary and extra-pulmonary TB) which were captured and implemented in the database during sequencing development in the lab ^38,39^. It should be noted that we excluded pleural TB as depending of the clinical presentation it can be considered as PTB or EPTB. Microbiological characteristics, such as Mtb lineage, smear results, time to positivity and drug resistance, were recorded (**Table S1**), as was patients’ demographic and baseline descriptive characteristics (**Table 1**). Regarding patients’ descriptive characteristics, only data available between 2 weeks before TB diagnosis and 1 week after initiation of anti-TB treatment or nutritional supplementation were considered.

### TB-associated severity indices

Three TB-associated severity indices were assessed in this study.

The modified Bandim score is the most largely used PTB prognosis score. It considers 5 symptoms (cough, hemoptysis, dyspnea, chest pain, night sweats) and 5 clinical findings (anemia, tachycardia, positive finding at lung auscultation, fever, BMI<18 and <16), with one point for each. One clinical finding was excluded, the mid upper arm circumference (MUAC) as this data was not available in the Lyon University Hospital. Accordingly, patients were stratified into two severity classes, mild (Bandim score ≤4) and moderate or severe (≥5) ^26,27,45^. The nutritional status of TB patients was evaluated thanks to the Malnutrition Universal Screening Tool (MUST), which include three variables: unintentional weight loss score (weight loss < 5% = 0, weight loss 5–10% = 1, weight loss > 10% = 2), BMI score (BMI>20.0 = 0, BMI 18.5–20.0 = 1, BMI<18.5 = 2) and anorexia (if yes = 2). Malnutrition is frequently observed in patients with PTB and a previous study showed a poorer prognosis for PTB patients with MUST ≥4 ^28^.

The nutritional risk score is a four-points score including both nutritional and immune characteristics: low BMI (<18.5), hypoalbuminaemia (<30.0 g/L), hypocholesterolaemia (<4mmol/L) and severe lymphocytopenia (<0.7 G cells/L). A high nutritional risk score (≥3) has been shown to be associated to poor prognosis in PTB ^29,46^.

### Culture of Mycobacterium tuberculosis

The routine laboratory diagnostic workflow consisted of treatment of pulmonary samples with the modified Kubica’s digestion-decontamination method ^47^, followed by inoculation in Mycobacteria Growth Indicator Tubes (MGITs) incubated in a BD BACTEC TM MGIT TM 960 instrument (BD, Sparks, MD, USA). Extrapulmonary samples were inoculated using the same medium without prior decontamination. Mtb genomic DNA extractions were performed after a single round of culture. Biobank Mtb isolates were inoculated in MGIT until exponential phase before Mtb genomic DNA extraction.

### Whole genome sequencing

Genomic DNA of Mtb positive cultures was purified from cleared lysate using a QIAamp DNA mini Kit (Qiagen). DNA libraries were prepared with Nextera XT kit (Illumina, San Diego, USA). Samples were sequenced on NextSeq or MiSeq system (Illumina) to produce 150 or 300 base-pair paired-end reads at the Bio-Genet NGS facility of Lyon University Hospital, as previously described ^38^. Reads were mapped with BOWTIE2 to the Mtb H37Rv reference genome (Genbank NC000962.2) and variant calling was made with SAMtools mpileup, as previously described ^38^.

### Illumina data analysis

A valid nucleotide variant was called if the position was covered by a depth of at least 10 reads and supported by a minimum threshold rate of 10%. Regions with repetitive or similar sequences were excluded, i.e. regions of PE, PPE, PKS, PPS, ESX genes. The reference genome coverage breadth was at least 93% with a mean depth of coverage of at least 50x.

### Variant assignment

In a previous study, we showed no significant differences in variant detection and frequencies between sequencing on direct samples and after subculture on media used in routine practice ^6^. Moreover, for this study, 10 isolates were extracted and sequenced twice to evaluate the variability in mutation frequencies between sequencing experiments. In both sequencing experiments, 52 unfixed mutations were detected at similar frequencies (±10%), ranging from 10 to 90% (**Fig. S9A**). Accordingly, to identify the minimum number of variant in each Mtb clinical isolate, a variant was defined as an assembly of mutations at frequencies of ±10% as illustrated in **Fig. S9B**.

### Mtb α-diversity indices

The alpha diversity index was calculated by using the software R statistic, Package (vegan) version 2.5-7.

### Selection analysis

We used PAML package to test for selection pressures in our dataset. This method uses powerful statistics to test for heterogeneous dn/ds ratios at different positions and/or in different branches. It has already been successfully used in MTC genome to confirm that positions involved in drug resistance and some positions in membrane proteins are subjected to positive selection ^48^. Shortly, this method allows to compare scenarios including (H1) or not (H0) higher dn/ds ratio on some residues. This means that two categories of sites are set up that have different distributions of their dn/ds (options can force some of the characteristics of these distributions, here both M1 and M2 models distributions were explored). The method allows to identify whether scenario H1 is more likely than H0 using a Likelihood Ratio Test (true if p-value<0.05).

Here we explored whether genes having the same functions (as characterized by available annotation) have a higher probability to be under selection in the evolutionary branches leading to one type of localization or the other (pulmonary versus extra-pulmonary). These tests were performed using the branch-site model of PAML.

While compiling the statistics enabling LRT tests described above, PAML package also associates a probability to each codon that underwent positive selection presure: the Bayesian Empirical Bayes (BEB) posterior probability that this codon has a dn/ds ratio higher than 1 as compared to the alternative scenario, starting from a (true if P>0.95).

To do so, we built the matrix including all SNPs for all variants (intra et inter patients variations). For each SNP, we reconstituted the corresponding amino-acid using annotation available from mycobrowser Release 2 (https://mycobrowser.epfl.ch/releases). The variant phylogeny was reconstructed using RAxML using a GTRCAT model. All terminal branches leading to pulmonary isolates were labelled for identifying selection in the lungs (“Pulmonary” analysis). All terminal branches leading to any extra-pulmonary isolate were labelled for identifying selection in microaerophilic organs *i.e.* organs other than the lungs (“Extra-Pulmonary” analysis).

### Statistical analysis

#### Univariate analysis

The study variables were expressed as count (percentage, %) for dichotomous variables and as medians (interquartile range [IQR]) for continuous values. The number of missing values was excluded from the denominator. Non-parametric statistical methods Fisher exact test, χ2 test, Mann-Whitney U test and Kruskal-Wallis analysis, using Dunn’s Multiple Comparison Test were used to compare groups, where appropriate. Statistical analyses were performed with Graph Pad Prism 5. *p<0.05, ** p <0.01, *** p <0.001.

#### Response variable and subsetting explanatory variables

We built a model to relate Mtb micro-diversity in pulmonary (P) and extra pulmonary (EP) isolates to various explanatory variables obtained for most patients in the cohort. The response variable we chose to model is a binary (0/1) measure of whether Mtb displayed or not some diversity as observed through counts of unique SNP profiles.

Among all the potential explanatory variables recorded in the initial dataset, we retained a chosen subset, following a series of filters:

1. We first removed 15 variables based on three non-exclusive conditions: (i) relatively low coverage in the cohort, (ii) little to no relationship with the response variable based on the literature, (iii) important redundancy with other variables in the dataset (e.g. percentage of neutrophil when the raw neutrophil count was also included);
2. Among the remaining variables, we removed all those that had less than 90% coverage (i.e. those that had missing values for 10% or more of the patients). Apart from the location of the infection (P vs. EP), the final consolidated dataset included 31 variables: 10 categorical variables and 21 quantitative ones. This dataset consisted of 282 rows (patient:TBlocation), 200 for pulmonary locations and 82 for EP locations. 42 patients had both microbiologically proven P and EP locations, so the final dataset included 240 patients (158 with only P location, 40 with only EP location, and 42 patients with both locations).

#### Model comparison

In a first exploratory phase, we looked for all models explaining the response variable using a limited number of explanatory variables. We used generalized linear modelling, assuming that the binary response variable could be modelled through a binomial distribution and using a logit transformation linking response to explanatory variables. With 31 possible explanatory variables, the number of potential models to test is very high and impossible to tackle (2^31^, i.e. more than 2 billion models). In order to reduce this complexity, we chose to restrict our search to models incorporating between 0 and 12 variables. As presented in the result, this approach was sufficient to obtain “best models” that had fewer than 12 variables, hence hinting at the uselessness of pursuing our search further into models of higher complexity.

To rank the different tested models, we used the Akaike Information Criterion corrected for small sample sizes (AICc) ^49^ because we expected many tapered effects of the different variables ^50^. Models with the lowest AICc values were the ones that had the best goodness-of-fit. Models were computed and compared using the ‘glmulti’ package version 1.0.8 in the R software version 3.6.3 ^51^. To optimize computation time, we first looked for all models with 0 – 9 variables among the 31 present in the dataset, and then looked for models with 10, 11 and 12 variables (thus, 4 parallel uses of glmulti). For each of these four chunks of model comparison, we retained only the 500 best models (sensu AICc) and gathered all these models in a consensus pool of good models. The importance of model variables was then assessed using Akaike weights of each of the 31 variables by summing the Akaike weights of all models incorporating the focal variable ^50,52^. Since each variable was incorporated in exactly half of the tested models, the importance is expected to be ½ for variables that did not modify model fit better than the null expectation. Following Massol et al. (2007)^52^, we thus considered that all variables that were *likely* to have an effect on the response variable were those with an importance larger than 1/(1+*e*^−1^), i.e. 0.73, *plausible* or *implausible* on either side of 0.5, and unlikely when variable importance was lower than 0.27. For prediction purposes, we retained all models that were within 2 units of the best model’s AICc and used multi-model inference based on this set of models ^50^. We computed unconditional variance using the method of Johnson and Omland (2004)^53^ and obtained confidence intervals on model predictions using the method suggested by Burnham and Anderson ^50^, using function ‘predict’ in the R package ‘glmulti’.

#### Parameter value comparisons

When categorical variables had an effect on the response variable, we tested for pairwise differences in the coefficients associated to the different levels of the categorical variables. To do so, we used the R package ‘emmeans’ version 1.4.5, which tested pairwise differences between marginal means averaged over all values of other categorical variables (e.g. differences between “Mtb lineages” were assessed by averaging the effect of “double location”), using asymptotic test on the z-score obtained from the pair of coefficients ^54^.

#### Estimating multi-model errors

Once the set of good models had been determined, we re-sampled the dataset in order to obtain estimates of the multi-model prediction errors (i.e. false negatives and positives). For a given target proportion of initial dataset rows to be included in the training set, we drew a random sample of rows stratified by unique combinations of categorical variables used by the multi-model, i.e. we made sure that all combinations of factors were included in the training set. This resulted in actual fraction of data rows in the training dataset slightly higher than the target proportion because some combinations of categorical variable modalities were quite rare and thus systematically added to the training set.

With a given training dataset, we considered the rest of the dataset as validation dataset. We fitted the whole set of good models using only the training dataset, with possibly estimates of model coefficients different from those obtained with the whole dataset due to sampling. Based on the probability of observing some Mtb micro-diversity for all samples in the training set, we looked for a threshold on this probability that would maximize the true skill statistic (TSS) if the multi-model were to predict diversity when model predictions were above this threshold and no diversity otherwise. TSS is a simple statistic (also called informedness or Youden’s J statistic) equal to the sum of sensitivity and specificity minus one. This threshold optimization procedure was performed using function ‘optim.thresh’ in R package ‘SDMTools’ version 1.1-221.2 ^55^.

The performance of the multi-model inferred from the training dataset was assessed by predicting the response variable in the validation dataset, using the above-mentioned threshold for predicting presence/absence of Mtb micro-diversity. This prediction yielded a confusion matrix (observed vs. predicted absence/presence of Mtb micro-diversity) which was then analysed using standard statistics, i.e. its sensitivity, specificity, accuracy (1 - error rate) and TSS.

## ACKNOWLEDGMENTS

This work was supported by the LABEX ECOFECT (ANR-11-LABX-0048) of Université de Lyon, within the program “Investissements d’Avenir” (ANR-11-IDEX-0007) operated by the French national research agency (Agence nationale de la recherché, ANR).

The funders had no role in study design, data collection and analysis, decision to publish, or preparation of the manuscript.

