## Supplementary Figures and Tables for "Within-host genetic micro-diversity of *Mycobacterium tuberculosis* and the link with tuberculosis disease features"

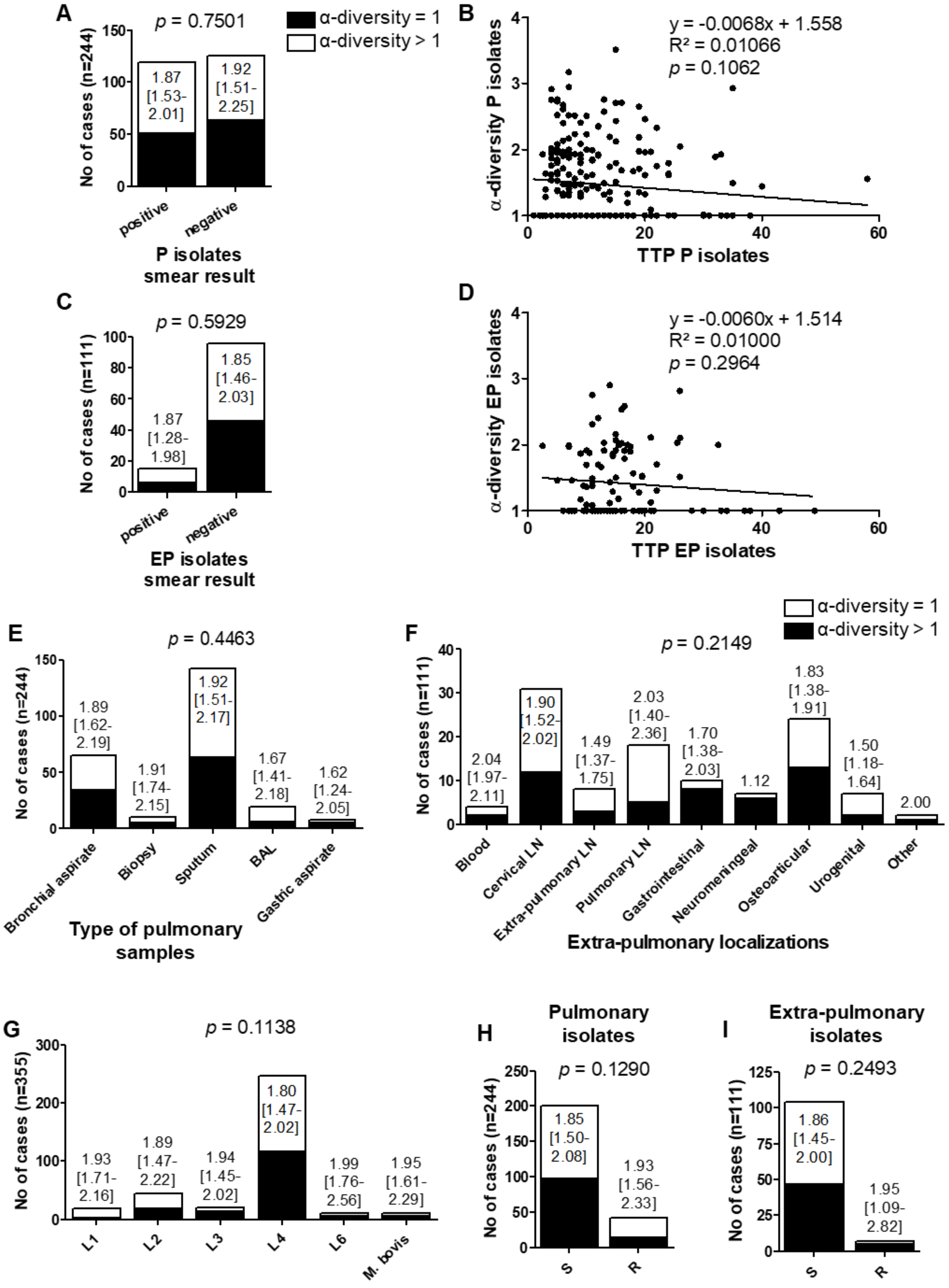

**Figure S1: No correlation between the detection of micro-diversity or the range of α-diversity in Mtb clinical isolates and Mtb characteristics.**

Association between the detection and the range of Mtb α-diversity and smear results (A and C), time to positivity of clinical isolates (B and D), type of pulmonary samples (E), localization of extra-pulmonary TB (F), Mtb lineage (G) and Mtb resistance status (H and I). P isolate: pulmonary isolate; EP isolate: extra-pulmonary isolate; TTP: time to positivity of culture sample; BAL: broncho-alveolar lavage; LN: lymph node; L1: lineage 1; S: antibiotic sensitive; R: resistant to at least one first-line anti-TB drug. A, C, E-I. Black bar: α-diversity=1 no diversity detected by WGS; white bar: α-diversity>1 at least two variants detected by WGS. *p* = x.xxxx: non-parametric statistical methods Fisher exact test or $\chi 2$ test were used to compare groups where appropriate. *p-value* < 0.05 was considered significant. x.xx [y.yy-z.zz]: median [IQR] of Mtb α-diversity. Mann-Whitney U test or Kruskal-Wallis analysis, using Dunn's Multiple Comparison Test, where appropriate, were used to compare ranges of Mtb α-diversity between groups (no statistical differences observed). B and D. Linear regression between time to positivity (TTP) of pulmonary (P) or extra-pulmonary (EP) clinical isolates and their respective Mtb α-diversities.

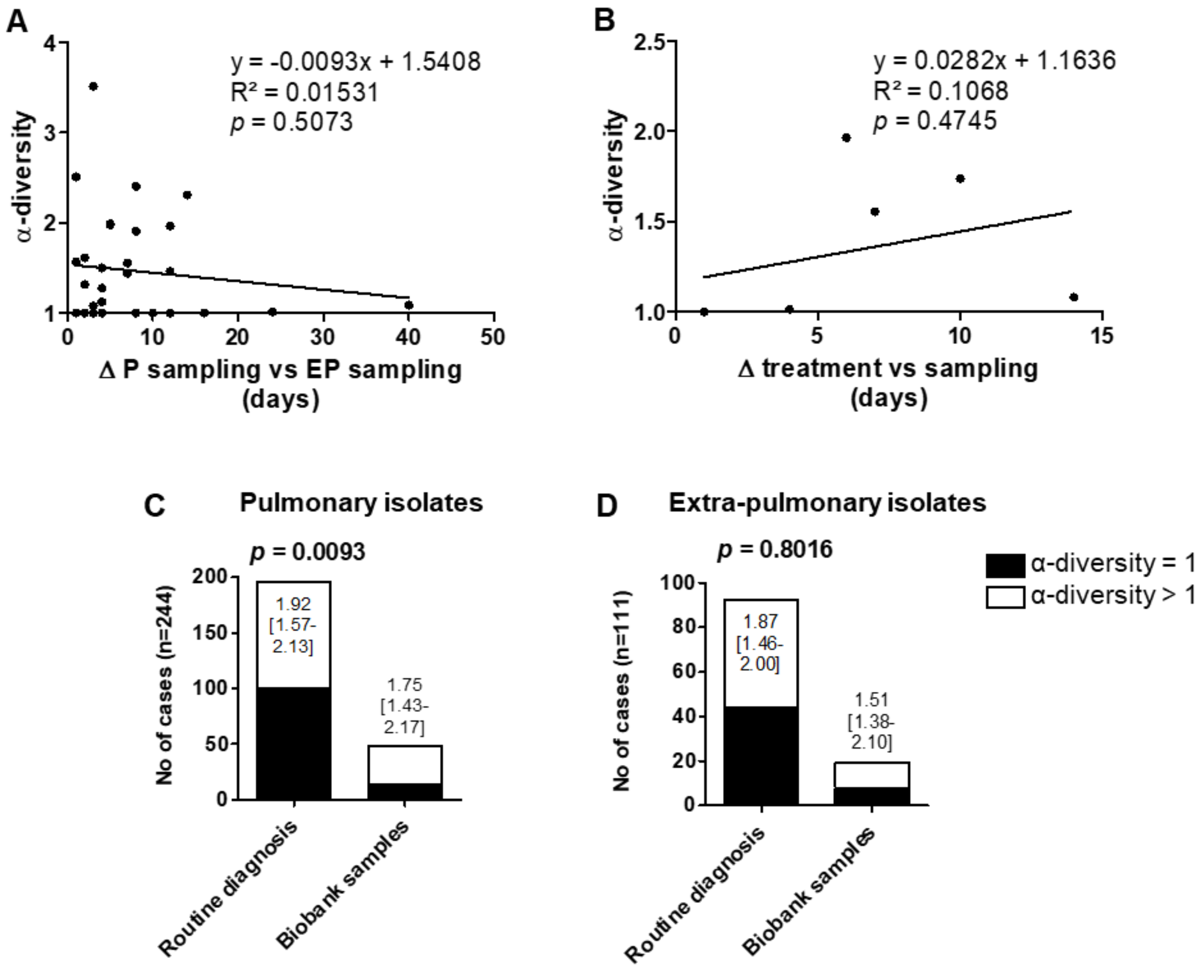


**Figure S2: Risks of bias due to Mtb sampling.**

(A and B) Linear regression between Mtb α-diversity and days between pulmonary and extra-pulmonary sampling (A) and days between treatment initiation and Mtb sampling (B), only concerning isolates from patients with both microbiologically proven PTB and EPTB. (C and D) Detection and range of Mtb α-diversity from pulmonary (C) and extra-pulmonary (D) samples analyzed by WGS after a single round of culture (routine diagnosis) or after a single freeze/thaw cycle (biobank samples). Black bar: α-diversity=1 no diversity detected by WGS; white bar: α-diversity>1 at least two variants detected by WGS. *p* = x.xxxx: non-parametric statistical method Fisher exact was used to compare groups. *p-value* < 0.05 was considered significant. x.xx [y.yy-z.zz]: median [IQR] of Mtb α-diversity. Mann-Whitney U test was used to compare ranges of Mtb α-diversity between groups (no statistical differences observed).

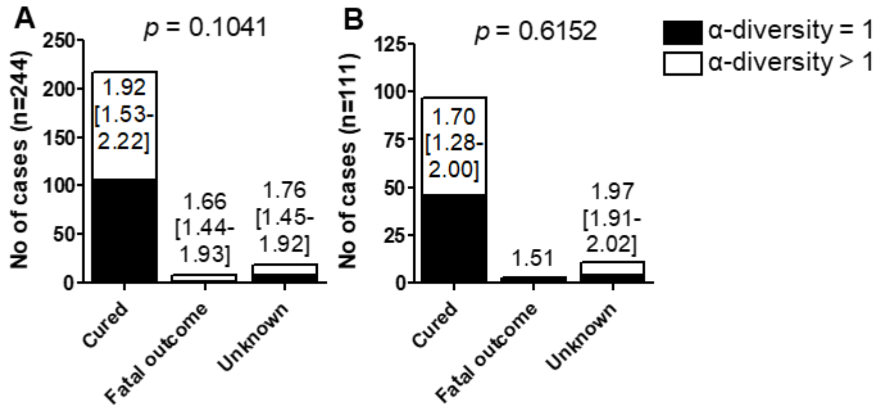


**Figure S3: Micro-diversity in Mtb clinical isolates and TB outcome**

Association between the detection and the range of Mtb α-diversity from pulmonary (A) and extra-pulmonary (B) samples and TB outcome. Unknown outcome: loss of follow-up or follow-up in another care facility. Black bar: α-diversity=1 no diversity detected by WGS; white bar: α-diversity>1 at least two variants detected by WGS. *p* = x.xxxx: non-parametric statistical methods $\chi 2$ test were used to compare groups. *p-value* < 0.05 was considered significant. x.xx [y.yy-z.zz]: median [IQR] of Mtb α-diversity. Kruskal-Wallis analysis, using Dunn's Multiple Comparison Test was used to compare ranges of Mtb α-diversity between groups (no statistical differences observed).

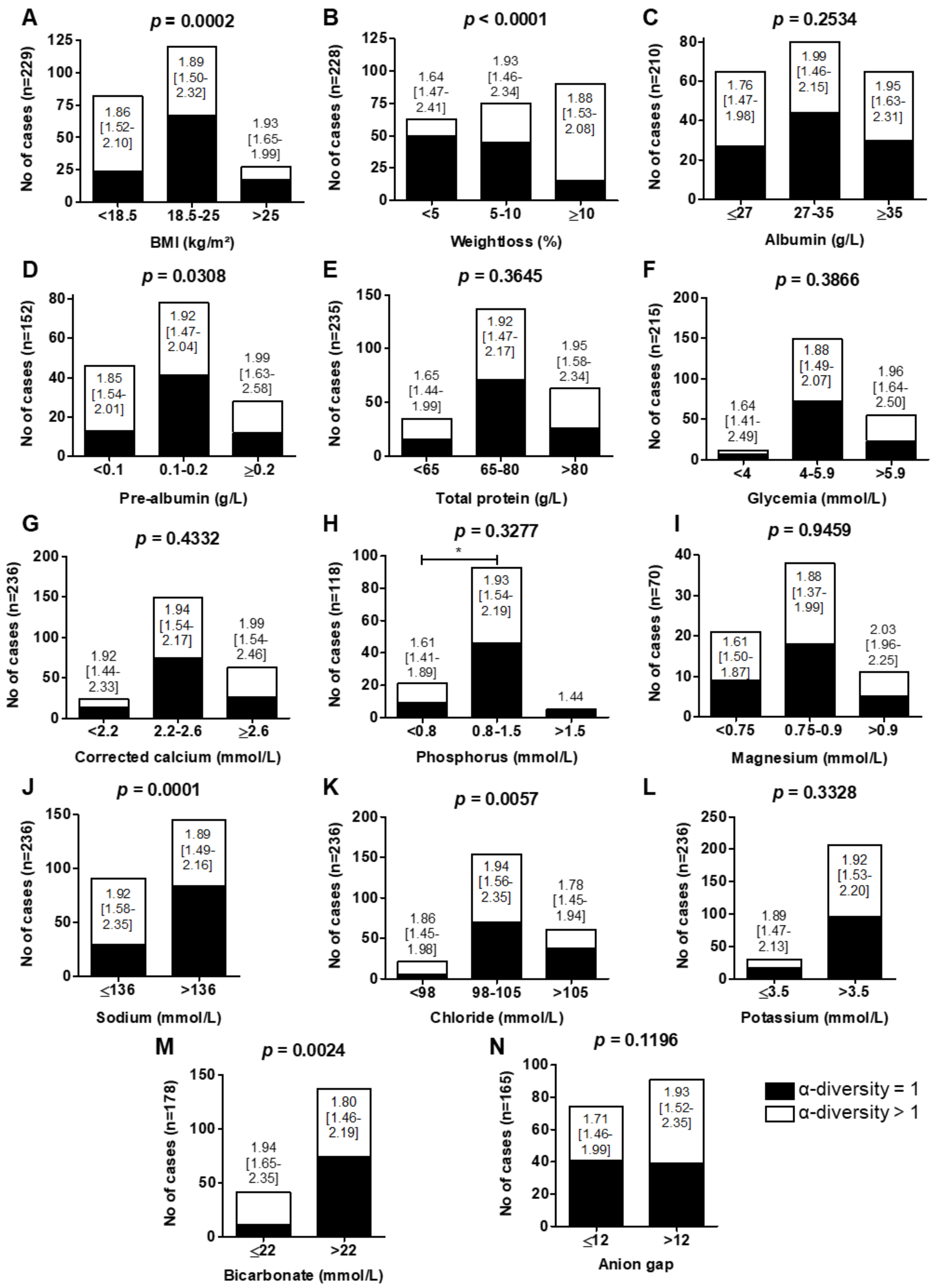


**Figure S4: The detection of genetic micro-diversity in pulmonary Mtb isolates is associated with malnutrition indices**

Association between the detection and the range of Mtb α-diversity from pulmonary samples and patient body mass index (BMI, A), unintentional weight loss (B), serum albumin (C), pre-albumin (D), total protein (E), glycemia (F), corrected calcium (G), phosphorus (H), magnesium (I), sodium (J), chloride (K), potassium (L), bicarbonate (M), and anion gap (N). Black bar: α-diversity=1 no diversity detected by WGS; white bar: α-diversity>1 at least two variants detected by WGS. $p$ = x.xxxx: non-parametric statistical methods Fisher exact test or $\chi 2$ test were used to compare groups where appropriate. *p-value* $< 0.05$ was considered significant. x.xx [y.yy-z.zz]: median [IQR] of Mtb α-diversity. Mann-Whitney U test or Kruskal-Wallis analysis, using Dunn's Multiple Comparison Test, where appropriate, were used to compare ranges of Mtb α-diversity between groups. * $p<0.05$.

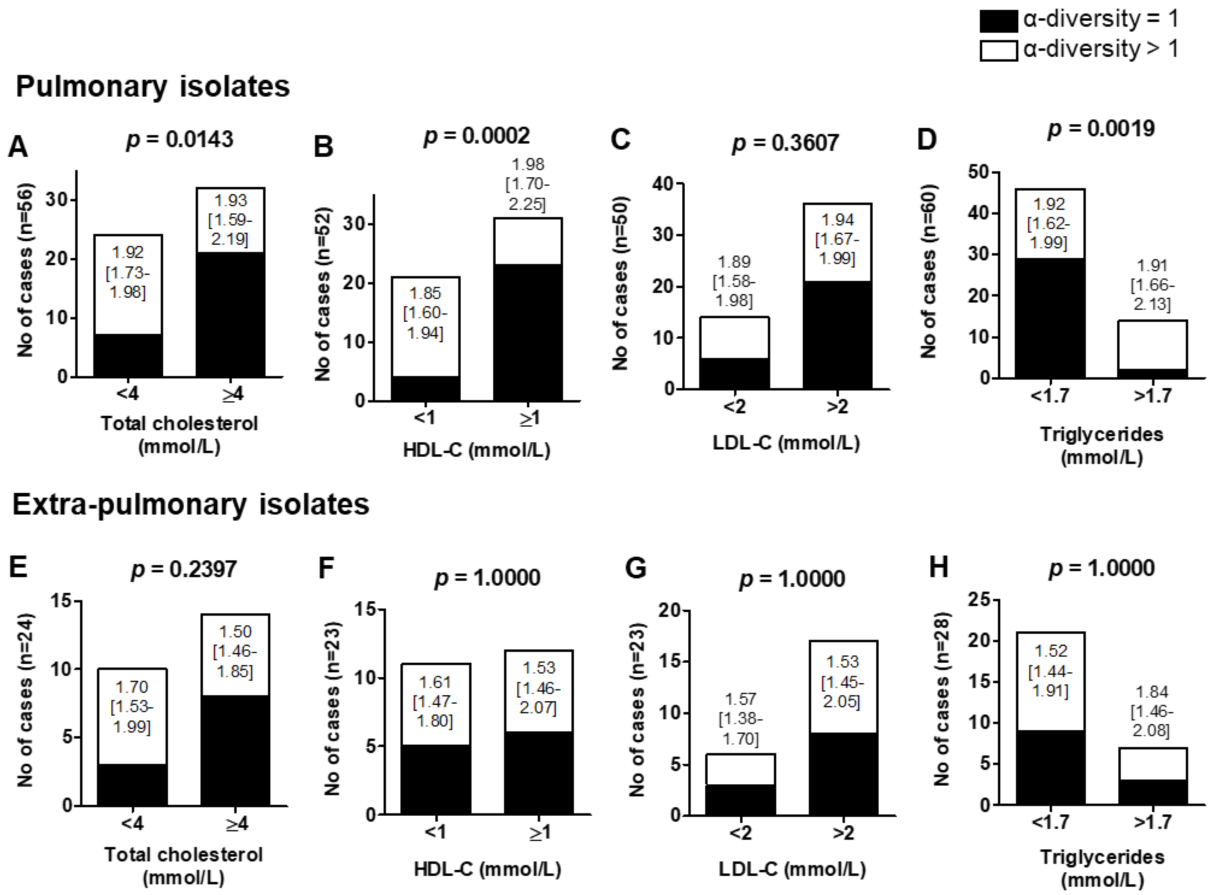


**Figure S5: Genetic micro-diversity in pulmonary and extra-pulmonary Mtb isolates and association with patients' serum lipid profile.**

Association between the detection and the range of Mtb α-diversity from pulmonary (A-D) and extra-pulmonary samples (E-H) and total cholesterol (A and E), HDL-C (B and F), LDL-C (C and G) and triglycerides (D and H). Black bar: α-diversity=1 no diversity detected by WGS; white bar: α-diversity>1 at least two variants detected by WGS. *p* = x.xxxx: non-parametric statistical methods Fisher exact test was used to compare groups. *p-value* $< 0.05$ was considered significant. x.xx [y.yy-z.zz]: median [IQR] of Mtb α-diversity. Mann-Whitney U test was used to compare ranges of Mtb α-diversity between groups (no statistical differences observed).

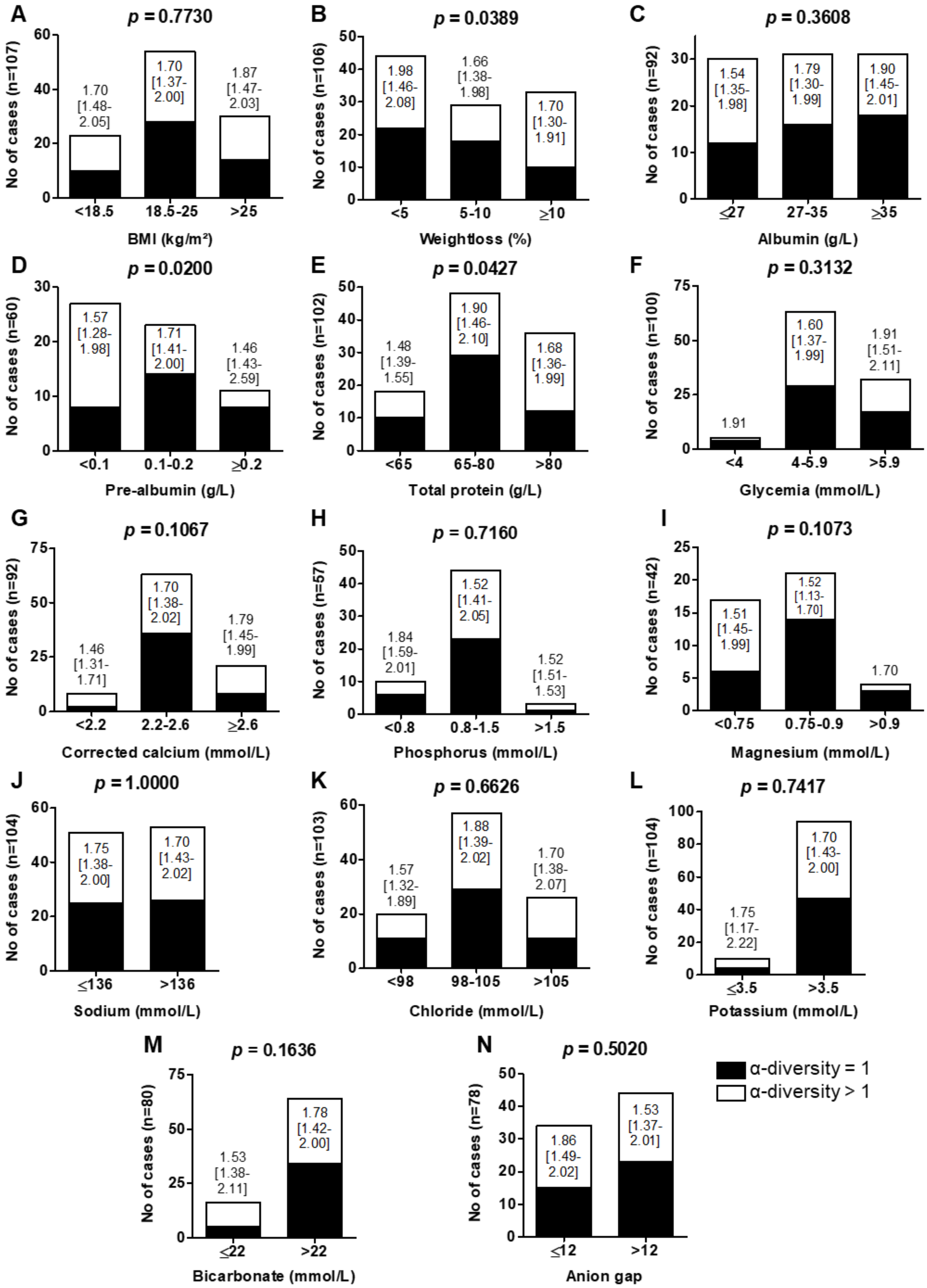


**Figure S6: The detection of genetic micro-diversity in extra-pulmonary Mtb isolates is marginally associated with malnutrition indices.**

Association between the detection and the range of Mtb α-diversity from extra-pulmonary samples and patient body mass index (BMI, A), unintentional weight loss (B), serum albumin (C), pre-albumin (D), total protein (E), glycemia (F), corrected calcium (G), phosphorus (H), magnesium (I), sodium (J), chloride (K), potassium (L), bicarbonate (M), and anion gap (N). Black bar: α-diversity=1 no diversity detected by WGS; white bar: α-diversity>1 at least two variants detected by WGS. $p$ = x.xxxx: non-parametric statistical methods Fisher exact test or $\chi 2$ test were used to compare groups where appropriate. *p-value* $< 0.05$ was considered significant. x.xx [y.yy-z.zz]: median [IQR] of Mtb α-diversity. Mann-Whitney U test or Kruskal-Wallis analysis, using Dunn's Multiple Comparison Test, where appropriate, were used to compare ranges of Mtb α-diversity between groups (no statistical differences observed).

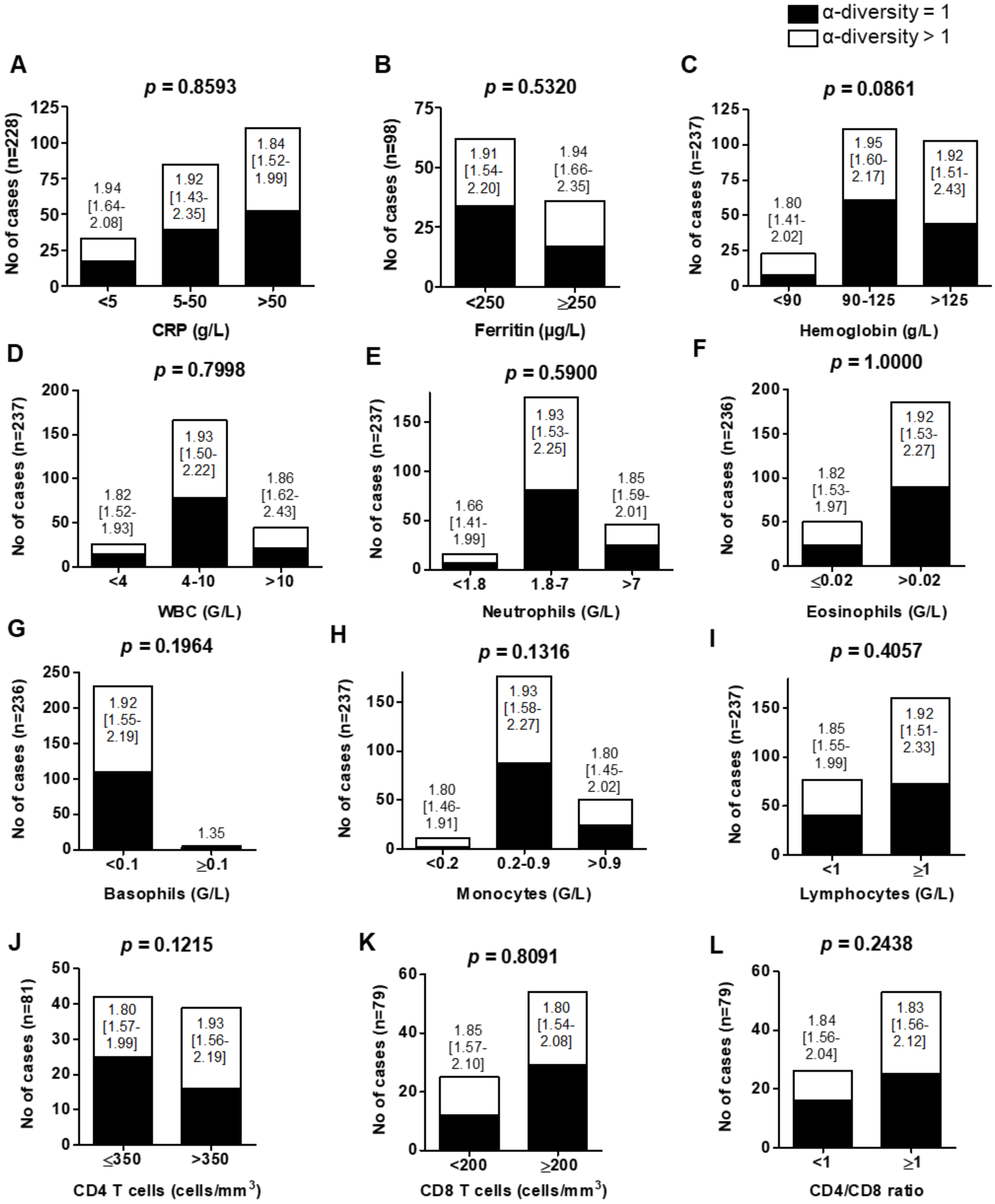


**Figure S7: The detection and the range of genetic micro-diversity in pulmonary Mtb isolates is not associated with inflammation or immune markers.**

Association between the detection and the range of Mtb α-diversity from pulmonary samples patient C-reactive protein (CRP, A), ferritin (B), hemoglobin (C) white blood cells (WBC, D), neutrophils (E), eosinophils (F), basophils (G), monocytes (H), lymphocyte (I), CD4 T cells (J), CD8 T cells (K) count in peripheral blood and CD4/CD8 ratio (L). Black bar: α-diversity=1 no diversity detected by WGS; white bar: α-diversity>1 at least two variants detected by WGS. *p* = x.xxxx: non-parametric statistical methods Fisher exact test or χ2 test were used to compare groups. *p-value* $< 0.05$ was considered significant. x.xx [y.yy-z.zz]: median [IQR] of Mtb α-diversity. Mann-Whitney U test or Kruskal-Wallis analysis, using Dunn's Multiple Comparison Test, where appropriate, were used to compare ranges of Mtb α-diversity between groups (no statistical differences observed).

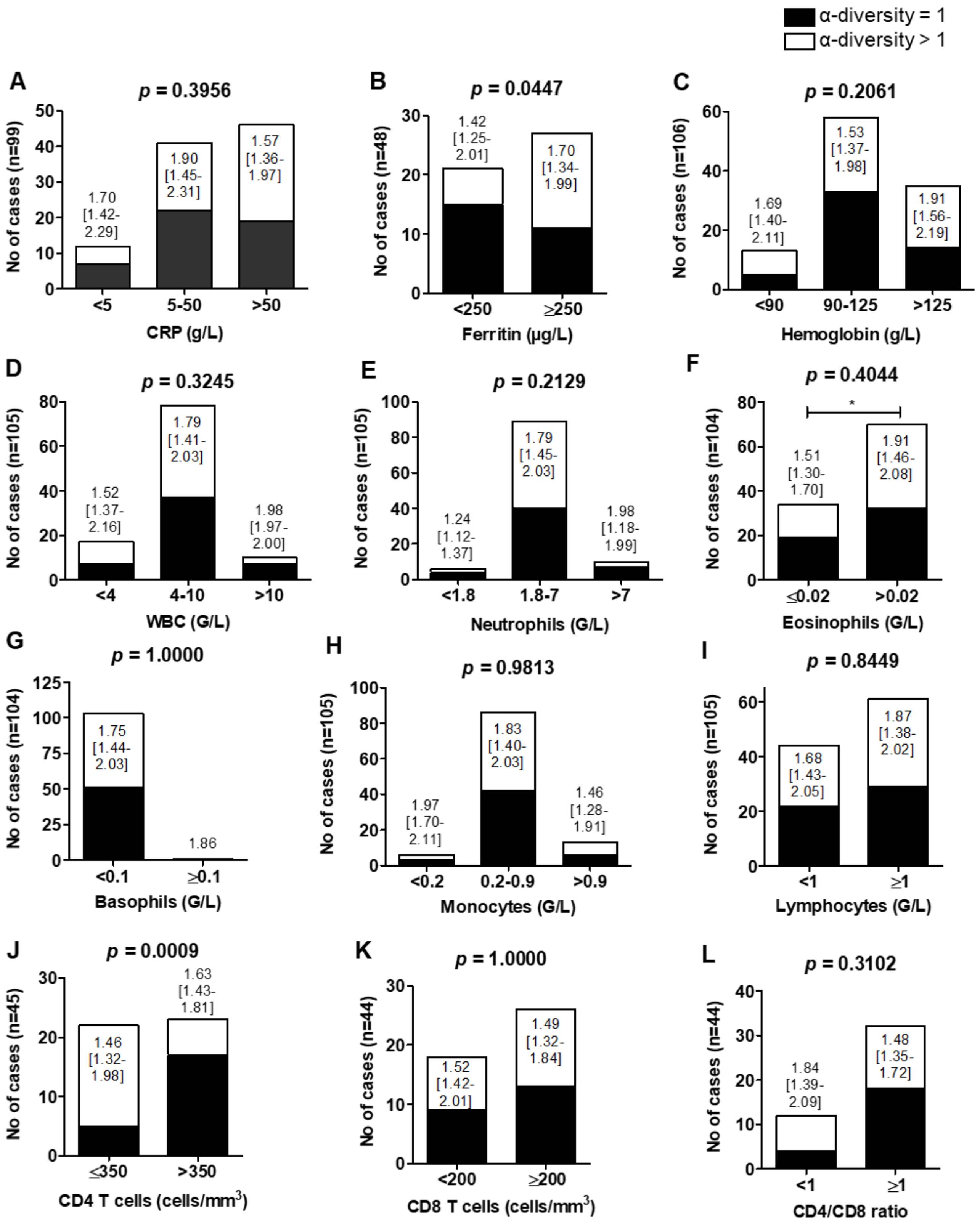


**Figure S8: The detection and the range of genetic micro-diversity in extra-pulmonary Mtb isolates is associated with high ferritin and low CD4 T cells count.**

Association between the detection and the range of Mtb α-diversity from extra-pulmonary samples patient C-reactive protein (CRP, A), ferritin (B), hemoglobin (C) white blood cells (WBC, D), neutrophils (E), eosinophils (F), basophils (G), monocytes (H), lymphocyte (I), CD4 T cells (J), CD8 T cells (K) count in peripheral blood and CD4/CD8 ratio (L). Black bar: α-diversity=1 no diversity detected by WGS; white bar: α-diversity>1 at least two variants detected by WGS. $p$ = x.xxxx: non-parametric statistical methods Fisher exact test or $\chi 2$ test were used to compare groups. *p-value* $< 0.05$ was considered significant. x.xx [y.yy-z.zz]: median [IQR] of Mtb α-diversity. Mann-Whitney U test or Kruskal-Wallis analysis, using Dunn's Multiple Comparison Test, where appropriate, were used to compare ranges of Mtb α-diversity between groups. * $p<0.05$.

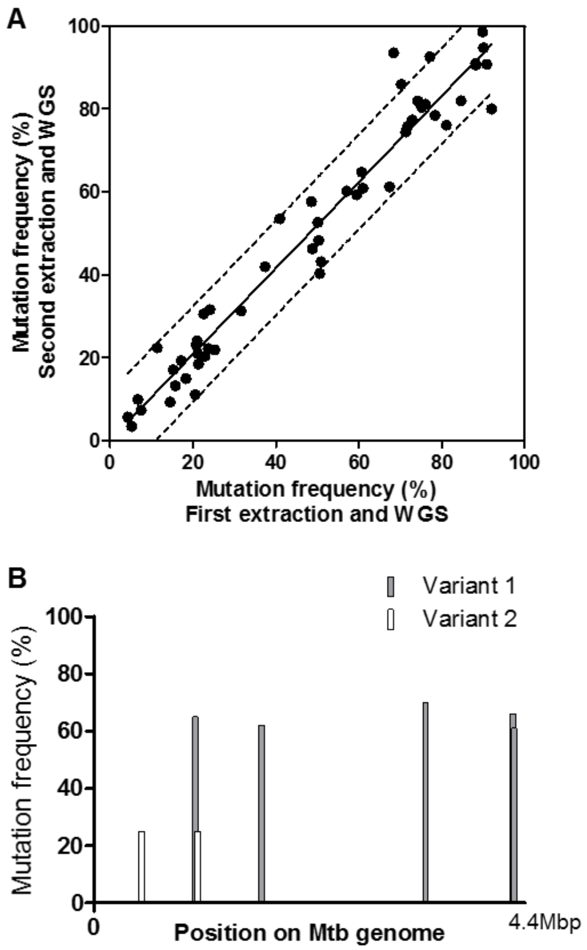


**Figure S9: Method for variant assignment in Mtb clinical isolates**

**A.** 10 isolates, containing 52 unfixed mutations at frequencies ranging from 10% to 90%, were extracted and sequenced twice to evaluate the variability in mutation frequencies between experiments. Continuous line is the linear regression curve. Discontinuous bands are the 90% prediction bands. **B.** Mutations frequencies across Mtb genome, obtained by WGS, each bar representing a mutation. For variant 1, 5 mutations at frequencies of 65, 62, 70, 66 and 61% were observed, indicating a frequency of 65% for this variant. For variant 2, 2 mutations were observed both at a frequency of 25%, indicating a frequency of 25% for this variant. A third variant, not carrying any of these 7 mutations, is estimated at a frequency of 10%.

**Table S1:** Microbiological characteristics of Mtb isolates included in the study

| **Microbiological characteristics** | **Pulmonary TB n=244** | **Extra-pulmonary TB n=111** | ***p-values*** |
|---|---|---|---|
| Type of P sample, n (%)<br>Bronchial aspiration<br>Biopsy<br>Sputum<br>BAL<br>Stomach tube | <br>65 (26.6)<br>10 (4.1)<br>142 (57.4)<br>19 (7.8)<br>8 (3.3) | N/A | N/A |
| EP localizations, n (%)<br>Blood<br>Cervical LN<br>Extra-pulmonary LN<br>Pulmonary LN<br>Gastrointestinal<br>Neuromeningeal<br>Osteoarticular<br>Urogenital<br>Cutaneous<br>Pericarditis | N/A | <br>4 (3.6)<br>31 (32.4)<br>8 (7.2)<br>18 (16.2)<br>10 (9.0)<br>7 (6.3)<br>24 (21.6)<br>7 (6.3)<br>1 (0.9)<br>1 (0.9) | N/A |
| Lineages, n (%)<br>L1<br>L2<br>L3<br>L4<br>L6<br>*M. bovis* | <br>10 (4.1)<br>33 (13.5)<br>14 (5.7)<br>145 (59.4)<br>5 (2.0)<br>7 (2.9) | <br>8 (7.2)<br>12 (10.8)<br>7 (6.3)<br>73 (65.8)<br>4 (3.6)<br>7 (6.3) | 0.3759 |
| Smear positive isolates, n (%) | 120 (49.1) | 15 (13.5) | **<0.0001** |
| TTP (days), median [IQR] | 10.0 [6-17] | 14.5 [11-19] | **<0.0001** |
| Resistant Mtb (to at least one first line anti-TB drug), n (%) | 43 (17.6) | 7 (6.3) | **0.0047** |
| α-diversity >1, n (%) | 130 (53.3) | 59 (53.2) | 1.0000 |
| α-diversity >1, median [IQR] | 1.88 [1.52-2.14] | 1.86 [1.45-2.00] | 0.2042 |
